## Supporting information for "Specificity of *Loxosceles* α clade phospholipase D enzymes for choline-containing lipids: role of a conserved aromatic cage"

**Table S1. Details of the simulation systems used. Each system was simulated for 300 ns (production run). Two replicas were performed for each system.**

| System | Protein | Lipid bilayer | Total number of atoms | Number of water molecules | Ions |
|---|---|---|---|---|---|
| 1 | Li_αIA1 | 256 POPC | 106131 | 22514 | 1 $Cl^-$<br>1 $Mg^{2+}$ |
| 2 | Ll_αIII1 | 256 POPC | 106757 | 22663 | 4 $Na^+$<br>1 $Mg^{2+}$ |
| 3 | St_βIB1 | 256 POPC | 114556 | 25277 | 3 $Na^+$<br>1 $Mg^{2+}$ |
| 4 | St_βIB1 R44Y/S60Y | 256 POPC | 106572 | 22613 | 4 $Na^+$<br>1 $Mg^{2+}$ |
| 5 | Li_αIA1 | 50 PSM<br>180 POPC<br>26 CHOL | 105028 | 22783 | 1 $Cl^-$<br>1 $Mg^{2+}$ |
| 6 | Ll_αIII1 | 50 PSM<br>180 POPC<br>26 CHOL | 105081 | 22741 | 4 $Na^+$<br>1 $Mg^{2+}$ |
| 7 | St_βIB1 | 50 PSM<br>180 POPC<br>26 CHOL | 112307 | 25164 | 3 $Na^+$<br>1 $Mg^{2+}$ |
| 8 | St_βIB1 | 128 POPC<br>128 POPE | 104812 | 22413 | 3 $Na^+$<br>1 $Mg^{2+}$ |

**Trajectory Analysis**

**Table S2. Inventory of the candidate atoms for the hydrophobic interaction analysis.** The atom names correspond to the nomenclature of the CHARMM36 force field.

| Residue/lipid | Candidate atoms |
|---|---|
| ALA | CA HA CB HB1 HB2 HB3 |
| ARG | CA HA CB HB1 HB2 CG HG1 HG2 |
| ASN | CA HA CB HB1 HB2 |
| ASP | CA HA CB HB1 HB2 |
| CYS | CA HA CB HB1 HB2 |
| GLN | CA HA CB HB1 HB2 CG HG1 HG2 |

| | |
|---|---|
| GLU | CA HA CB HB1 HB2 CG HG1 HG2 |
| GLY | CA HA1 HA2 |
| HSD | CA HA CB HB1 HB2 |
| HSE | CA HA CB HB1 HB2 |
| HSP | CA HA CB HB1 HB2 CG |
| ILE | CA HA CB HB CG1 HG11 HG12 CG2 HG21 HG22 HG23 CD HD1 HD2 HD3 |
| LEU | CA HA CB HB1 HB2 CG HG CD1 HD11 HD12 HD13 CD2 HD21 HD22 HD23 |
| LYS | CA HA CB HB1 HB2 CG HG1 HG2 CD HD1 HD2 CE HE1 HE2 |
| MET | CA HA CB HB1 HB2 CG HG1 HG2 CE HE1 HE2 HE3 |
| PHE | CA HA CB HB1 HB2 CG CD1 HD1 CD2 HD2 CE1 HE1 CE2 HE2 CZ HZ |
| PRO | CA HA CB HB1 HB2 CD HD1 HD2 CG HG1 HG2 |
| SER | CA HA CB HB1 HB2 |
| THR | CA HA CB HB CG2 HG21 HG22 HG23 |
| TRP | CA HA CB HB1 HB2 CG CD1 HD1 CD2 CE3 HE3 CZ3 HZ3 CH2 HH2 CZ2 HZ2 |
| TYR | CA HA CB HB1 HB2 CG CD1 HD1 CD2 HD2 CE1 HE1 |
| POPC/POPE | C23 H3R H3S C24 H4R H4S C25 H5R H5S C26 H6R H6S C27 H7R H7S C28 H8R H8S C29 H91 C210 H101 C211 H11R H11S C212 H12R H12S C213 H13R H13S C214 H14R H14S C215 H15R H15S C216 H16R H16S C217 H17R H17S C218 H18R H18S H18T C33 H3X H3Y C34 H4X H4Y C35 H5X H5Y C36 H6X H6Y C37 H7X H7Y C38 H8X H8Y C39 H9X H9Y C310 H10X H10Y C311 H11X H11Y C312 H12X H12Y C313 H13X H13Y C314 H14X H14Y C315 H15X H15Y C316 H16X H16Y H16Z |
| DOPC/DOPE | C12 C11 H11A H11B C1 HA HB C2 HS C22 H2R H2S C3 HX HY C32 H2X H2Y C23 H3R H3S C24 H4R H4S C25 H5R H5S C26 H6R H6S C27 H7R H7S C28 H8R H8S C29 C210 C211 H11R H11S C212 H12R H12S C213 H13R H13S C214 H14R H14S C215 H15R H15S C216 H16R H16S C217 H17R H17S C218 H18R H18S H18T C33 H3X H3Y C34 H4X H4Y C35 H5X H5Y C36 H6X H6Y C37 H7X H7Y C38 H8X H8Y C39 C310 C311 H11X H11Y C312 H12X H12Y C313 H13X H13Y C314 H14X H14Y C315 H15X H15Y C316 H16X H16Y C317 H17X H17Y C318 H18X H18Y H18Z |
| PSM | C4S H4S C5S H5S C6S H6S H6T C7S H7S H7T C8S H8S H8T C9S H9S H9T C10S H10S H10T C11S H11S H11T C12S H12S H12T C13S H13S H13T C14S H14S H14T C15S H15S H15T C16S H16S H16T C17S H17S H17T C18S H18S H18T H18U C3F H3F H3G C4F H4F H4G C5F H5F H5G C6F H6F H6G C7F H7F H7G C8F H8F H8G C9F H9F H9G C10F H10F H10G C11F H11F H11G C12F |

| | |
|---|---|
| | H12F H12G C13F H13F H13G C14F H14F H14G C15F H15F H15G C16F H16F H16G H16H |
| CHOL | C4 H4A H4B C5 C6 H6 C7 H7A H7B C8 H8 C14 H14 C15 H15A H15B C16 H16A H16B C17 H17 C13 C18 H18A H18B H18C C12 H12A H12B C11 H11A H11B C9 H9 C10 C19 H19A H19B H19C C1 H1A H1B C2 H2A H2B C20 H20 C21 H21A H21B H21C C22 H22A H22B C23 H23A H23B C24 H24A H24B C25 H25 C26 H26A H26B H26C C27 H27A H27B H27C |

**Figure S1. Evolution of the protein backbone Root Mean Square Deviation (RMSD) during all simulations involving a pure POPC bilayer.** Replica 1 is represented in orange and replica 2 in cyan.

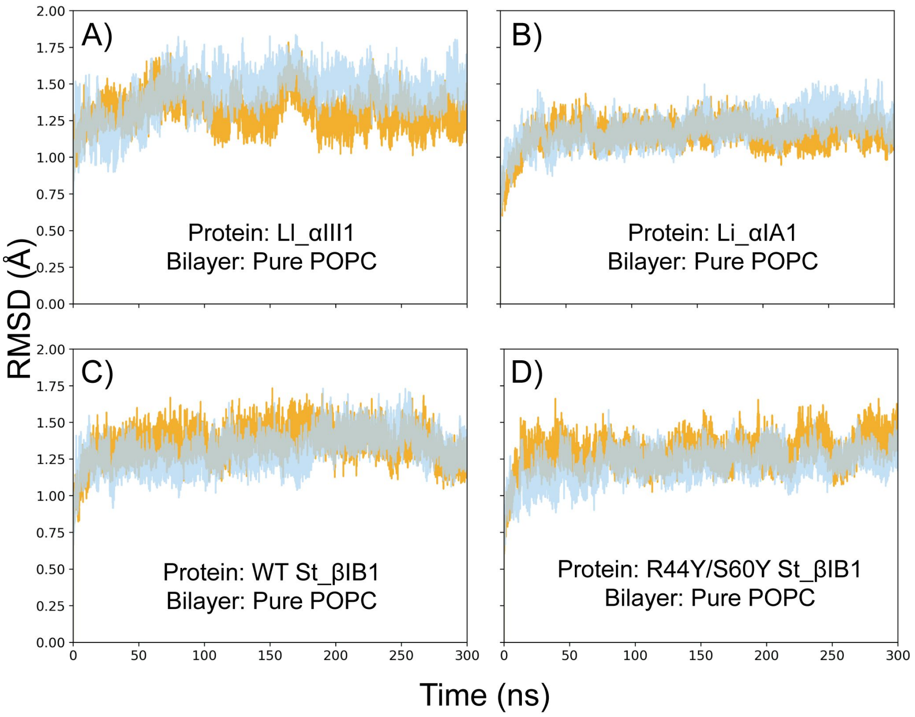

**Figure S2. Evolution of the protein backbone Root Mean Square Deviation (RMSD) during all simulations involving a PC:SM:CHOL (70:20:10) or a POPC:POPE (50:50).** Replica 1 is represented in orange and replica 2 in cyan.

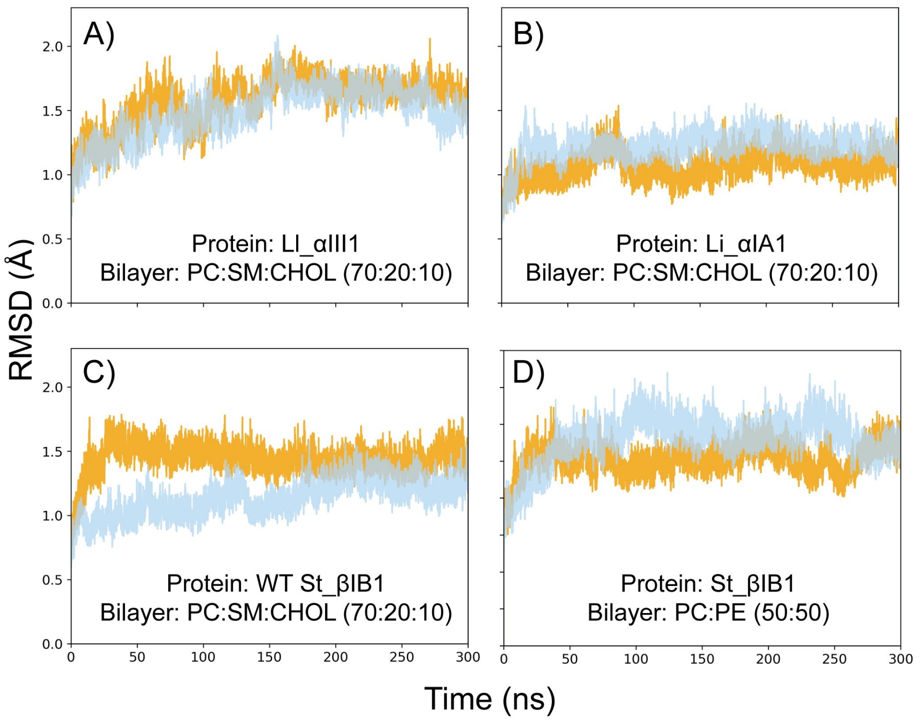


**Figure S3. Simulation of Li_αIA1, Ll_αIII1 and St_βIB1 on a pure POPC:PSM:CHOL (70:20:10) bilayer.** Snapshot at 0, 80, 160, 270 and 300 ns were extracted to follow the evolution of the systems during the simulations.

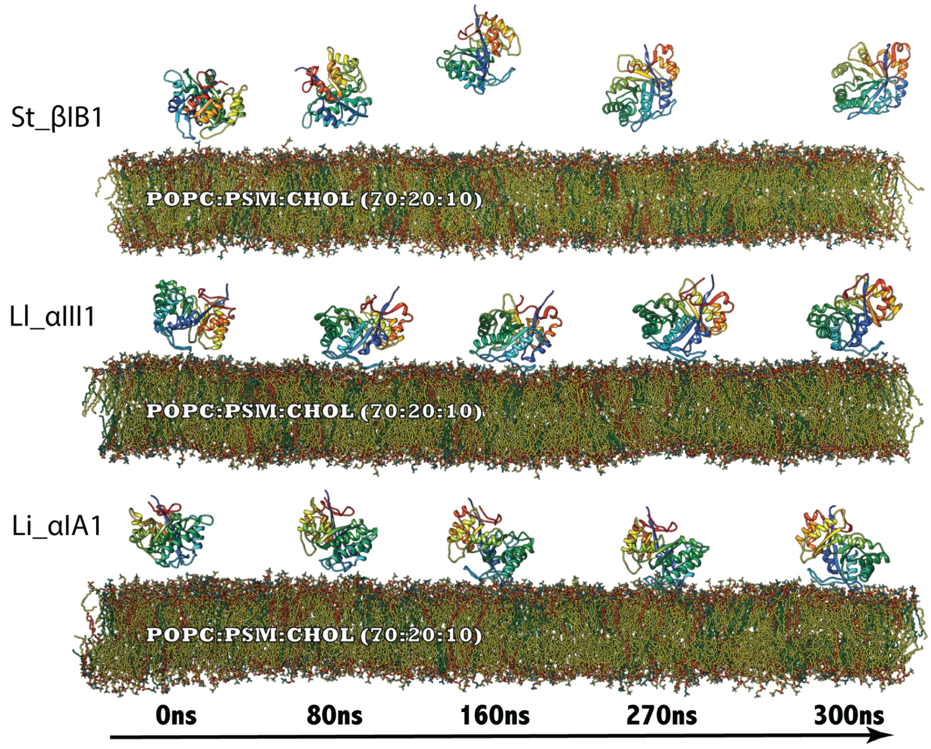

**Table S3. Anchoring depth of Li_αIA1, Ll_αIII1 and R44Y/S60Y St_βIB1 on a pure POPC bilayer.** The depth represents the distance between the C-alpha atom of each residue and the upper phosphate plane. R1 and R2 stand for replica 1 and replica 2. Only residues with a relevant anchoring depth are indicated in this table.

| SSE | Residue | Depth (Å) | | | | | |
|---|---|---|---|---|---|---|---|
| | | Li_αIA1 | | Ll_αIII1 | | R44Y/S60Y St_βIB1 | |
| | | R1 | R2 | R1 | R2 | R1 | R2 |
| β2α2 | 37 | 4.96 | 6.64 | 4.46 | 4.06 | 2.22 | 4.23 |
| | 38 | 2.39 | 3.93 | 1.26 | 0.64 | 0.79 | 2.56 |
| | 39 | 2.21 | 4.62 | 2.29 | 1.33 | 1.74 | 2.35 |
| | 44 | 5.45 | 6.70 | 3.64 | 3.24 | 5.87 | 4.43 |
| | 46 | 5.39 | 6.41 | 3.71 | 3.76 | 6.09 | 5.24 |
| | 48 | 1.56 | 2.28 | 0.45 | 0.80 | 6.11 | 2.62 |
| | 49 | -0.94 | -0.09 | -0.16 | 0.29 | 0.60 | 0.90 |
| | 50 | -0.50 | 0.21 | 0.88 | 1.67 | -1.01 | 0.34 |
| | 51 | 2.93 | 3.73 | 3.43 | 3.56 | 1.80 | 3.88 |
| | 52 | 4.90 | 6.02 | 5.67 | 5.03 | 3.61 | 6.12 |
| | 53 | 3.39 | 5.10 | 4.38 | 3.47 | 3.54 | 8.53 |
| | 54 | 3.97 | 5.88 | 5.09 | 5.68 | 0.39 | 7.06 |
| | 55 | 4.83 | 5.48 | 4.49 | 5.64 | 1.82 | 8.13 |
| | 56 | 2.71 | 3.21 | 1.66 | 3.02 | 2.33 | 6.76 |
| | 57 | 2.22 | 2.60 | 2.25 | 2.48 | 4.67 | 6.05 |
| | 58 | 1.54 | 1.78 | -0.79 | -0.39 | 2.47 | 3.49 |
| | 59 | 3.76 | 5.05 | 0.91 | 1.61 | 4.10 | 5.68 |
| | 60 | 3.80 | 5.80 | 1.95 | 2.42 | 5.33 | 5.31 |
| β3α3 | 95 | 2.52 | 5.09 | 4.85 | 4.93 | 5.24 | 2.13 |
| | 96 | 3.83 | 6.62 | 2.51 | 1.42 | 2.01 | 1.67 |
| | 97 | 3.95 | 7.06 | 4.88 | 2.34 | 3.75 | 2.56 |
| | 98 | 3.77 | 7.55 | 5.36 | 1.50 | 3.15 | 4.13 |
| β6α6 | 201 | 4.20 | 7.80 | 9.96 | 12.33 | 11.53 | 15.17 |
| | 202 | 4.40 | 7.94 | 12.80 | 15.34 | 13.49 | 15.64 |

**Table S4. Anchoring depth of Li_αIA1 and Ll_αIII1 on a POPC:PSM:CHOL (70:20:10) bilayer.** The depth represents the distance between the C-alpha atom of each residue and the upper phosphate plane. R1 and R2 stand for replica 1 and replica 2. Only residues with a relevant anchoring depth are indicated in this table.

| SSE | Residue | Depth (Å) | | | |
|---|---|---|---|---|---|
| | | Li_αIA1 | | Ll_αIII1 | |
| | | R1 | R2 | R1 | R2 |
| β2α2 | 37 | 7.36 | 7.04 | 7.18 | 5.29 |
| | 38 | 5.22 | 5.07 | 4.72 | 2.44 |
| | 39 | 4.76 | 4.25 | 5.22 | 2.14 |
| | 44 | 9.16 | 9.49 | 6.32 | 5.47 |
| | 46 | 7.50 | 6.96 | 4.53 | 4.89 |
| | 48 | 2.13 | 1.95 | 2.45 | 1.35 |
| | 49 | -0.38 | -0.48 | 0.18 | -0.30 |
| | 50 | -0.13 | -0.50 | 0.91 | 0.53 |
| | 51 | 3.46 | 2.32 | 4.08 | 3.78 |
| | 52 | 5.29 | 4.43 | 5.79 | 5.68 |
| | 53 | 4.05 | 3.44 | 3.88 | 4.53 |
| | 54 | 5.43 | 4.74 | 4.99 | 6.30 |
| | 55 | 6.28 | 5.45 | 5.49 | 5.87 |
| | 56 | 4.99 | 3.68 | 3.17 | 3.01 |
| | 57 | 4.21 | 4.09 | 2.44 | 2.25 |
| | 58 | 3.73 | 2.29 | 0.58 | 0.38 |
| | 59 | 7.36 | 5.46 | 3.71 | 3.14 |
| | 60 | 8.51 | 7.00 | 5.17 | 4.18 |
| β3α3 | 95 | 3.01 | 2.99 | 6.35 | 3.18 |
| | 96 | 1.68 | 2.15 | 4.14 | 1.73 |
| | 97 | 3.42 | 3.56 | 6.37 | 3.76 |
| | 98 | 3.43 | 4.11 | 6.28 | 2.58 |
| β6α6 | 201 | 3.30 | 3.02 | 8.79 | 8.10 |
| | 202 | 3.36 | 4.78 | 7.40 | 10.73 |

**Table S5. Anchoring depth of St_βIB1 on a POPC:POPE (50:50) bilayer.** The depth represents the distance between the C-alpha atom of each residue and the upper phosphate plane. R1 and R2 stand for replica 1 and replica 2. Only residues with a relevant anchoring depth are indicated in this table.

| SSE | residue | Depth | |
|---|---|---|---|
| | | R1 | R2 |
| β2α2 | H47 | 10.73 | 10.96 |
| | G48 | 9.98 | 9.98 |
| | V49 | 6.77 | 7.86 |
| | P50 | 4.42 | 6.72 |
| | C51 | 4.39 | 5.09 |
| | D52 | 3.66 | 4.51 |
| | C53 | 0.05 | 0.88 |
| | F54 | -1.84 | -1.30 |
| | R55 | 0.91 | 1.24 |
| | S56 | 2.35 | 2.46 |
| | C57 | 4.93 | 4.71 |
| | T58 | 6.39 | 5.66 |
| | R59 | 9.66 | 7.67 |
| β6 | D192 | 11.92 | 13.64 |
| | G193 | 8.75 | 10.49 |
| | I194 | 5.96 | 7.69 |
| β6α6 | T195 | 4.16 | 5.37 |
| | N196 | 2.68 | 3.25 |
| | C197 | -0.88 | -0.16 |
| | L198 | -1.00 | -0.60 |
| | P199 | 0.76 | 0.63 |
| | R200 | 3.85 | 4.98 |
| | D201 | 4.30 | 6.29 |
| | D202 | 7.40 | 9.20 |
| α6 | N203 | 8.63 | 11.24 |
| | R204 | 11.70 | 13.75 |
| β7α7 | W226 | 10.75 | 11.03 |
| | S227 | 8.59 | 8.49 |
| | I228 | 6.45 | 6.11 |
| | D229 | 5.54 | 3.41 |
| | K230 | 2.83 | 1.89 |
| α7 | E231 | 4.27 | 3.16 |
| | S232 | 2.8 | 2.53 |
| | S233 | 4.30 | 4.40 |
| | I234 | 7.61 | 7.39 |
| | E235 | 7.75 | 7.14 |
| | N236 | 6.99 | 7.48 |
| | A237 | 9.89 | 10.59 |

**Figure S4. Distances between the aromatic cage and the choline group of POPC 23, 63 and 82 during R1 simulation of Li_αIA1 on a pure POPC bilayer.** The distance reported here is the distance between the center of mass of the aromatic cycles involved in the aromatic cage and the nitrogen atom of a given lipid (POPC 23, POPC 63 or POPC 82).

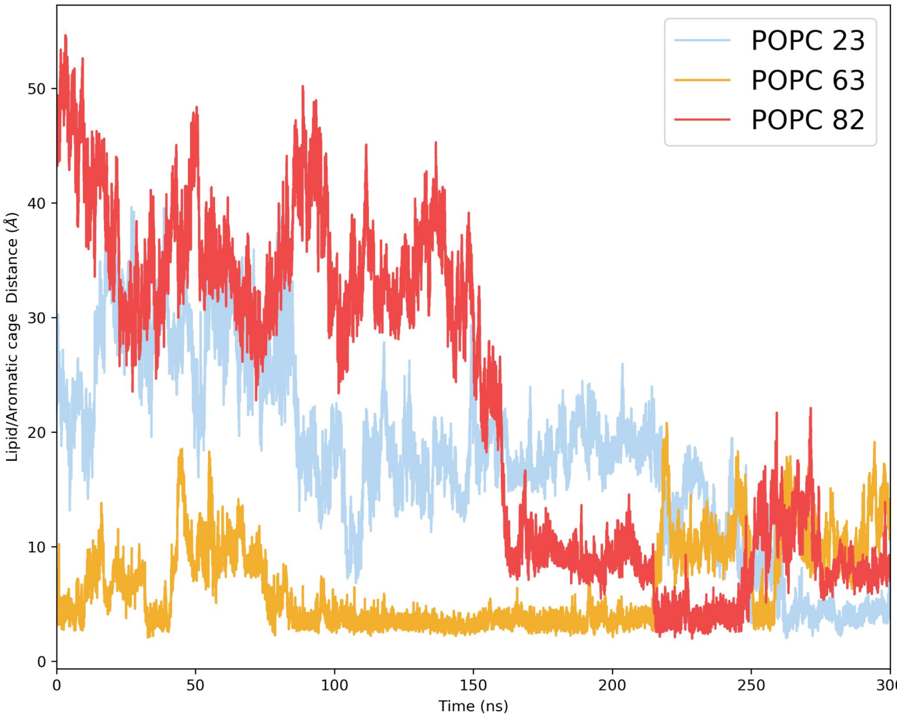


**Figure S5. Distance between the aromatic cage and the choline group of POPC 12 during R1 simulation of Ll_αIII1 on a pure POPC bilayer.** The distance reported here is the distance between the center of mass of the aromatic cycles involved in the aromatic cage and the nitrogen atom of POPC 12.

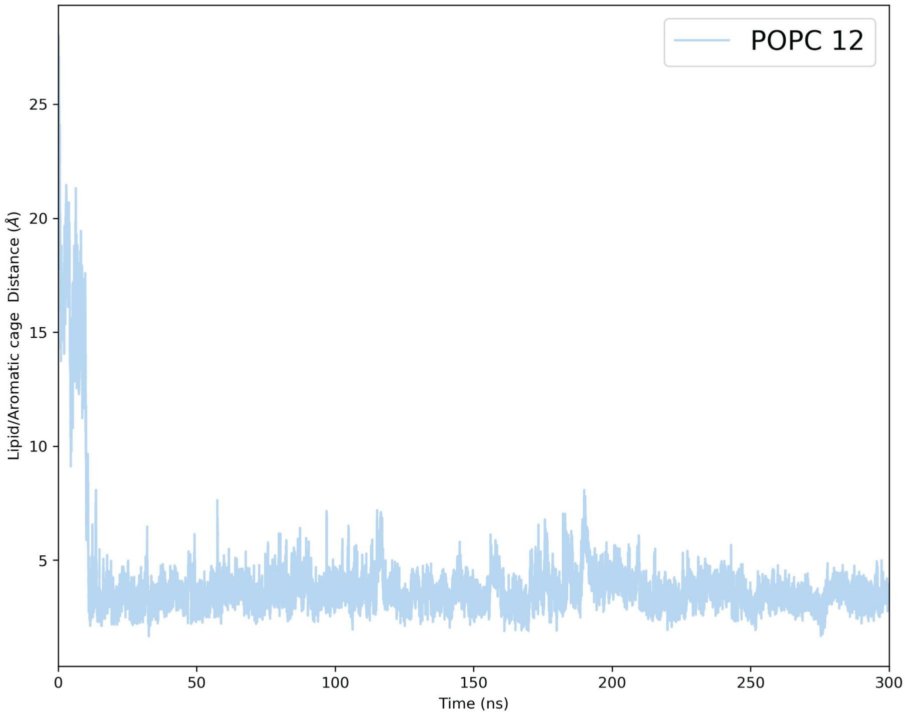

**Figure S6. Distances between the aromatic cage and the choline group of POPC 124 and 126 during R1 simulation of R44Y/S60Y St_βIB1 on a pure POPC bilayer.** The distance reported here is the distance between the center of mass of the aromatic cycles involved in the aromatic cage and the nitrogen atom of POPC 124 or POPC 126.

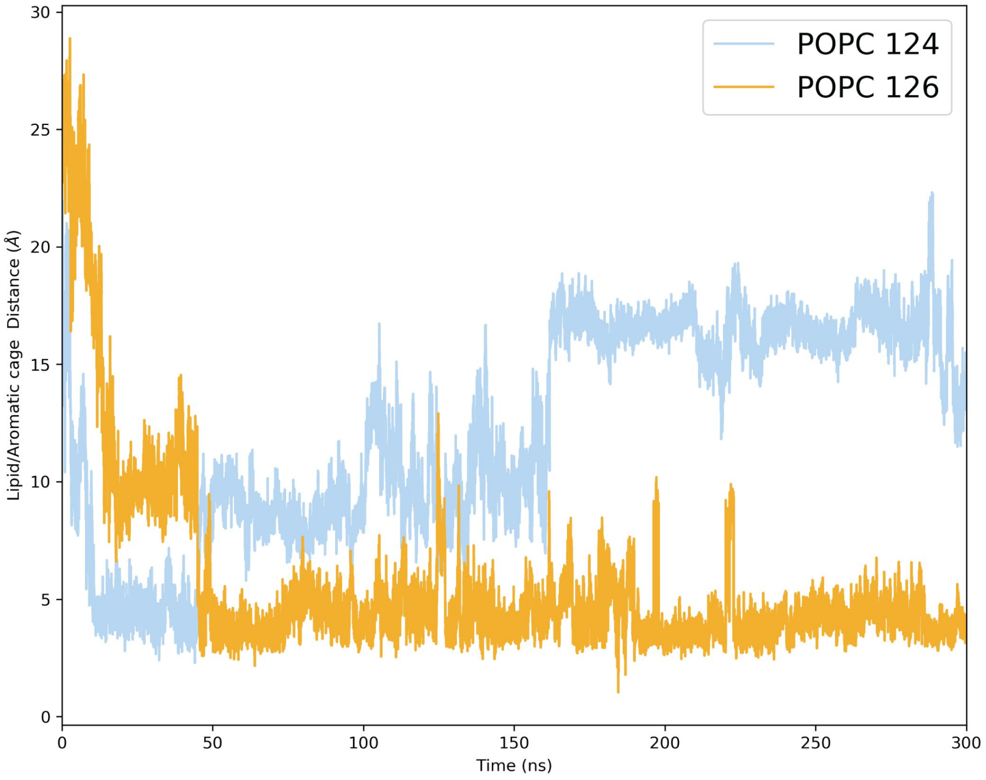


**Figure S7. Structural alignment between WT St_βIB1 and R44Y/S60Y St_βIB1.** The structures used for the alignment are final frames of each production run of WT St_βIB1 on a pure POPC bilayer and R44Y/S60Y St_βIB1 on a pure POPC bilayer. The calculated RMSD of the protein backbones is 0.99 Å.

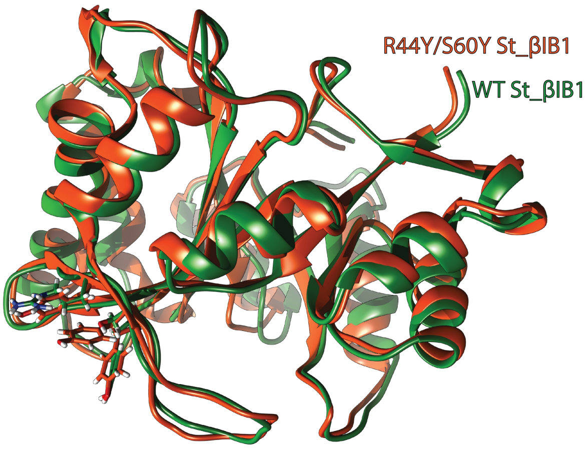

**Figure S8. Full multiple sequences alignment of α and β clade SicTox PLDs.** The sequences used for the generated the alignment are indicated in the main text. The part of the alignment with a green background represent the catalytic and the flexible loop. The protein simulated in this study are indicated with a yellow star. The red numbering corresponds to the numbering of the St_βIB1 structure (PDBid: 4Q6X). The black numbering indicates the position in the multiple sequence alignment.

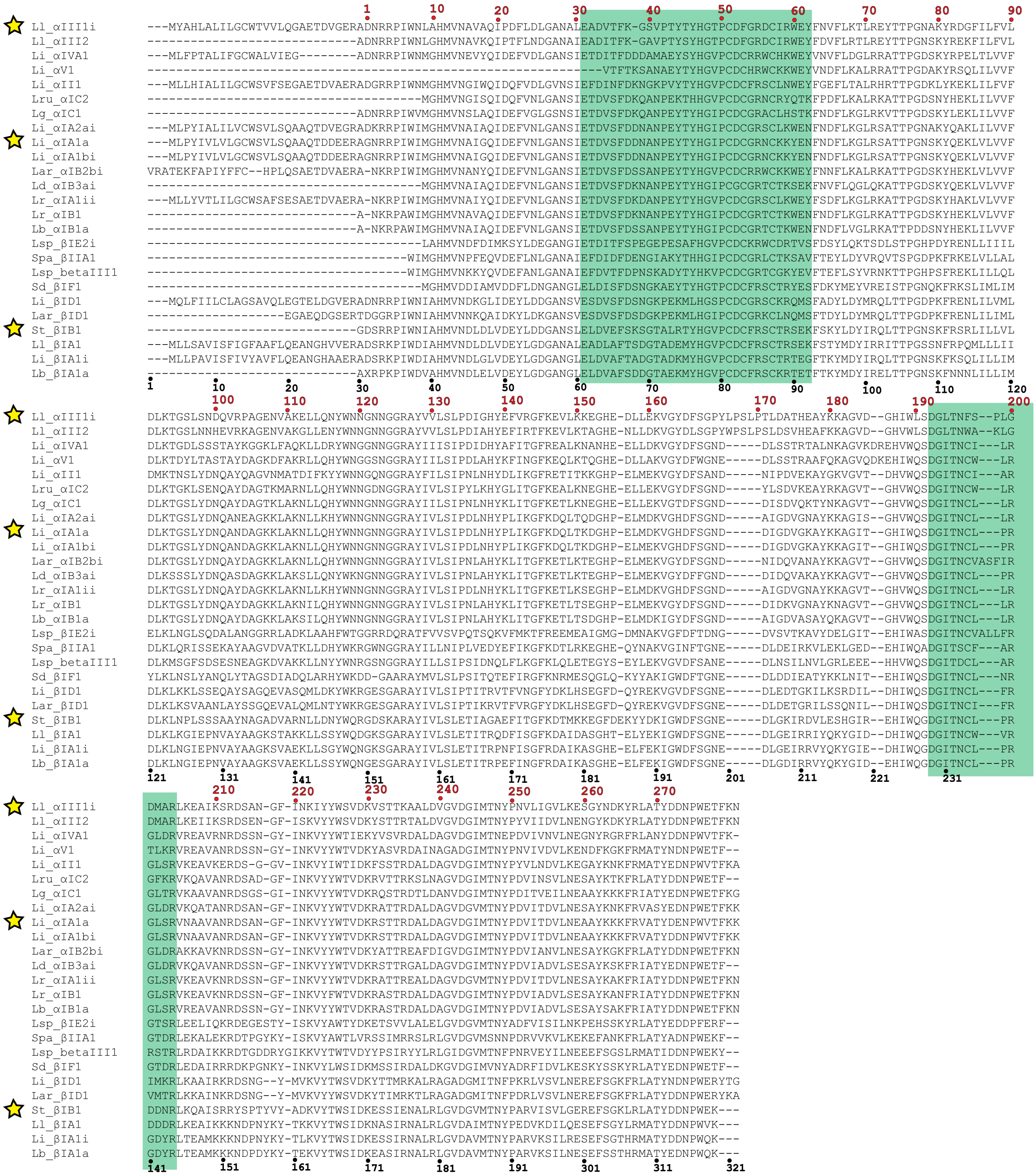
